# Chimera: Ultrafast and Memory-efficient Database Construction for High-Accuracy Taxonomic Classification in the Age of Expanding Genomic Data

**DOI:** 10.1101/2025.03.26.645388

**Authors:** Qinzhong Tian, Pinglu Zhang, Yanming Wei, Quan Zou, Yansu Wang, Ximei Luo

**Author notes:** These authors contributed equally to this work (Qinzhong Tian, Pinglu Zhang).

## Abstract

The rapid growth of genomic data expands species diversity but also causes taxonomic imbalance, with certain species heavily overrepresented. Both data volume and imbalance challenge the accuracy and efficiency of metagenomic tools. Here, we present Chimera, a transformative tool harnessing the Interleaved Merged Cuckoo Filter (IMCF) and FairMin-Cap (FMC) strategy for next-level performance. It achieves the highest classification accuracy while providing an astonishing 162-fold faster database assembly than Kraken2, constructing the complete RefSeq genome database within mere minutes using under 32 GB of RAM, enabling rapid and cost-effective database updates. Furthermore, Chimera’s universal memory scalability supports at least 300,000 species and potentially over 800,000 species in practical 1 TB systems, overwhelming traditional software solutions. Our results establish Chimera as a foundational tool for the next era of metagenomic research, laying a crucial cornerstone for the future of ultramassive genome datasets.

## 1. Background

Rapid advancements in sequencing technologies have led to exponential growth in metagenomic datasets, significantly enhancing our understanding of microbial ecology, clinical diagnostics, and biotechnology [1–3]. However, this rapid data expansion poses unprecedented challenges for metagenomic classification tools, primarily due to the increasing scale and complexity of reference databases [4–6]. Over the past decade, the NCBI RefSeq database has accumulated over 315,000 bacterial and archaeal genome assemblies, expanding by more than 35,000 genomes annually [7]. Similarly, the Genome Taxonomy Database (GTDB) has grown by over 270% since 2017, with an ongoing annual growth projection of approximately 30% [8]. Specialized platforms such as EMBL-EBI’s MGnify and the DOE JGI’s IMG/M have also experienced substantial data growth, further intensifying database complexity [9,10].

Existing metagenomic classification tools, including Kraken2, ganon, and Centrifuge, face significant limitations when managing extensive genomic databases containing hundreds of thousands or even millions of sequences [11–13]. These tools often require days to construct databases, and their runtime memory requirements frequently reach hundreds of gigabytes, preventing researchers from utilizing complete datasets effectively [5,14].

Additionally, imbalanced species representation within databases substantially reduces classification accuracy, a phenomenon known as Taxonomic Overrepresentation [15–18]. For instance, in 2017, twenty pathogenic bacterial species accounted for more than half of the prokaryotic genomes included in RefSeq, and they continue to represent a significant proportion today [19]. Such Taxonomic Overrepresentation obscures critical signals from less abundant or underrepresented taxa, emphasizing the urgent need for tools capable of efficiently maintaining balanced species representation in regularly updated databases [16,20].

To overcome these limitations, we introduce Chimera, a novel metagenomic classification tool specifically optimized for efficient database construction and accurate microbial identification. Chimera integrates two key innovations: the Interleaved Merged Cuckoo Filter (IMCF) and the FairMin-Cap (FMC) strategy. The IMCF significantly enhances query performance and reduces false positives, ensuring high classification accuracy. Concurrently, FMC addresses species overrepresentation by limiting minimizer counts per species, thus reducing database redundancy and memory usage dramatically. Notably, FMC’s universal applicability allows integration into other metagenomic classification frameworks, enhancing their database efficiency, representation balance, and classification accuracy. Furthermore, Chimera leverages Single Instruction, Multiple Data (SIMD) technology to further accelerate classification speed and throughput.

Experimental validations demonstrate Chimera’s superior performance in database construction speed and accuracy. Chimera constructs the complete RefSeq genome database approximately 162 times faster than Kraken2 and 74 times faster than Taxor, achieving completion within about five minutes using less than 32 GB of memory. Such extraordinary efficiency enables rapid and cost-effective database updates. Remarkably, Chimera is uniquely capable of building the entire RefSeq database within a 1TB memory constraint. In practical applications, Chimera requires approximately 1.2 MB of storage per species under typical configurations, enabling the theoretical accommodation of over 800,000 species within 1TB of memory. This advancement significantly alleviates dependency on high-performance computing resources, making high-quality classification analyses accessible even on standard personal computers. Future applications of Chimera to even larger and more complex metagenomic datasets will further demonstrate its role as a cornerstone for the upcoming era of ultramassive genome sets, transforming large-scale microbial classification from computationally intensive to routine.

## 2. Results

### 2.1 Utilizing Chimera in Metagenomic Taxonomic Classification

In large-scale metagenomic analysis, existing classification tools commonly suffer from prolonged database construction times, high memory requirements, and inconsistent classification performance. To overcome these challenges, we developed Chimera, a highly efficient metagenomic classification tool characterized by exceptionally fast database construction, minimal memory usage, and superior classification accuracy.

Chimera provides a streamlined, automated workflow that seamlessly handles the entire process—from downloading datasets from NCBI RefSeq to database construction—while enabling subsequent abundance analyses and interactive visualization using Krona (Figure 1) [21,22]. Chimera achieves its remarkable efficiency by integrating two key innovations: the IMCF and the FMC strategy. IMCF employs an interleaved design akin to interleaved Bloom filters, allowing multiple cuckoo filters to be queried simultaneously while retaining rapid query performance [23,24]. Because each cuckoo filter stores a 16-bit fingerprint— requiring only a single placement per item—it typically exhibits lower false-positive rates and faster construction compared to Bloom filters. However, to mitigate the increased space usage from storing 16 bits per entry, IMCF develops a merged approach in which the first 4 bits index the species and the remaining 12 bits encode the fingerprint. This design enables a single cuckoo filter to accommodate up to 16 species, substantially improving overall memory efficiency. Meanwhile, FMC provides a comprehensive approach to database optimization by filtering low-frequency minimizers, removing redundancies, and capping minimizer counts per species, thereby reducing taxonomic overrepresentation and enhancing classification accuracy.

**Figure 1.**
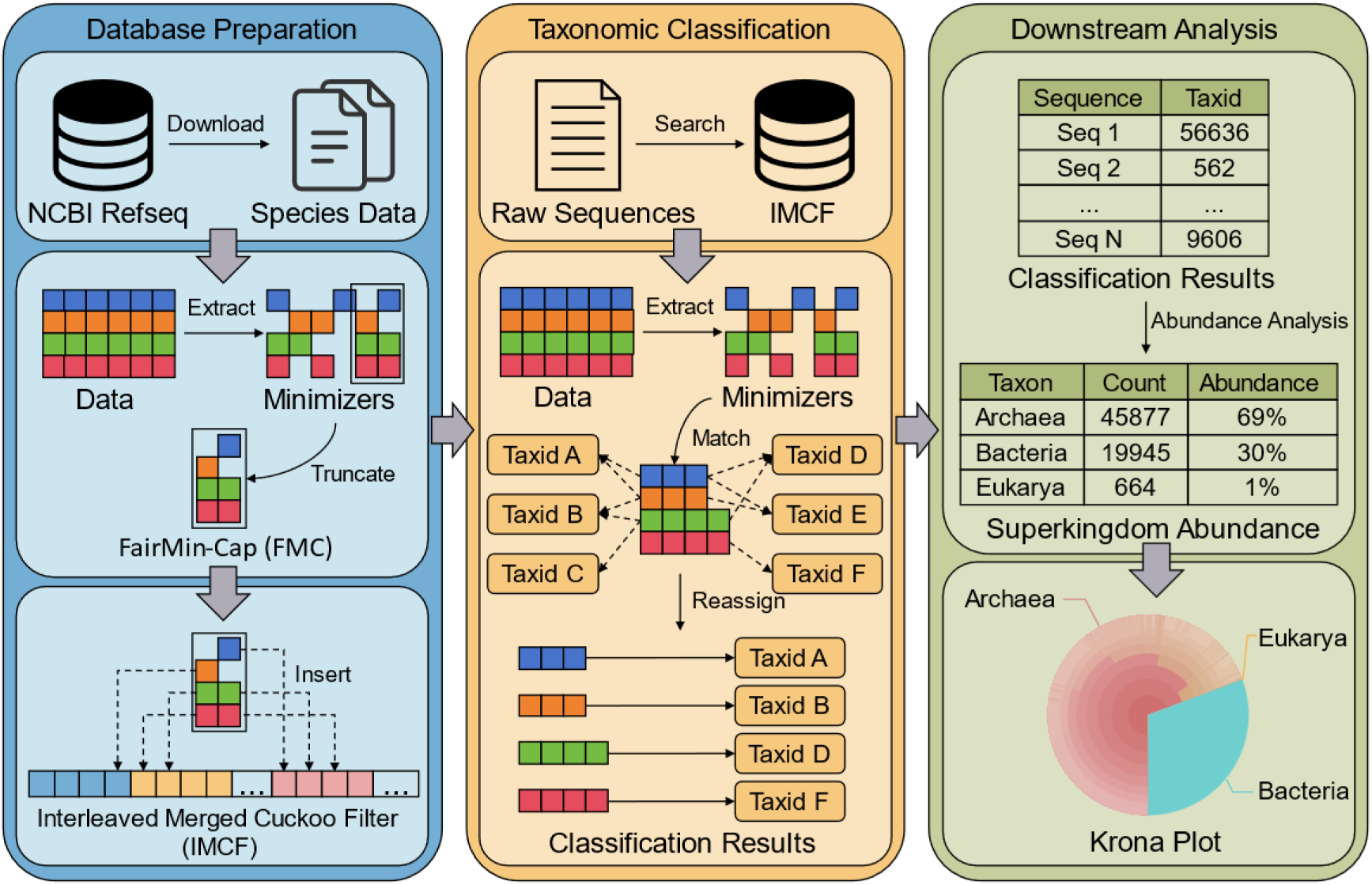
Workflow of Chimera for Metagenomic Taxonomic Classification. This figure illustrates the workflow of Chimera for metagenomic taxonomic classification, which consists of three stages: database preparation, taxonomic classification, and downstream analysis. In the database preparation stage, Chimera automatically downloads reference genome sequences from NCBI RefSeq, converts the sequences into Minimizers, and applies FMC for truncation optimization. The optimized Minimizers are then inserted into the IMCF to construct the database. In the taxonomic classification stage, Raw Sequences are processed by extracting Minimizers and querying them in the IMCF. The matching results are then refined using the Expectation-Maximization (EM) algorithm for reassignment, ultimately producing the final Classification Results. In the downstream analysis stage, Abundance Analysis is performed based on the classification results, and the figure illustrates an example at the Superkingdom level, producing Abundance Results, which are then visualized using Krona, where Archaea, Bacteria, and Eukarya are represented in different colors.

Furthermore, Chimera’s classification phase incorporates multiple optimizations, including a four-step filtering procedure, SIMD acceleration, and a flexible taxonomic assignment strategy that supports Variational Expectation-Maximization (VEM), Expectation-Maximization (EM), or Lowest Common Ancestor (LCA), with VEM as the default. These enhancements collectively ensure high-speed, accurate taxonomic classification across diverse metagenomic datasets.

The following sections comprehensively evaluate Chimera’s performance across three critical dimensions: database construction efficiency, classification accuracy, and the effectiveness of the FMC strategy.

### 2.2 Comparison of Database Construction Efficiency Across Tools

To comprehensively assess Chimera’s efficiency in database construction, we benchmarked its performance against five widely used metagenomic classification tools: Kraken2, Bracken, Ganon, Ganon2, and Taxor [11,12,14,25,26]. Kraken2 and Bracken are widely employed for metagenome analysis, whereas Ganon, Ganon2, Taxor, and Chimera all employ different variants of Bloom filters to optimize database storage and query efficiency.

We evaluated the tools across four datasets of varying scale and complexity: the smallest-scale Archaea database, the Complete RefSeq genome database (Complete), a reduced RefSeq database containing one genome assembly per species (CompleteONE), and the full RefSeq database. These datasets cover a broad range of taxonomic complexities, ensuring comprehensive and representative evaluations (Supplemental Table S1). Within Chimera, the load factor represents the proportion of utilized space within the IMCF, effectively defining how densely each cuckoo filter is populated. In all datasets except Archaea, we applied the default load factor settings to maintain optimal balance between memory efficiency and query performance. However, due to the simplicity and small size of the Archaea dataset, we configured it with an exceptionally high load factor of 0.95 to maximize space utilization. All experiments were conducted under identical hardware conditions—an AMD EPYC 7763 CPU with 1 TB memory, uniformly employing 32 computational threads. Key performance metrics, including construction time, peak memory usage, and database size, are reported in Figure 2 and Supplemental Table S2.

**Figure 2.**
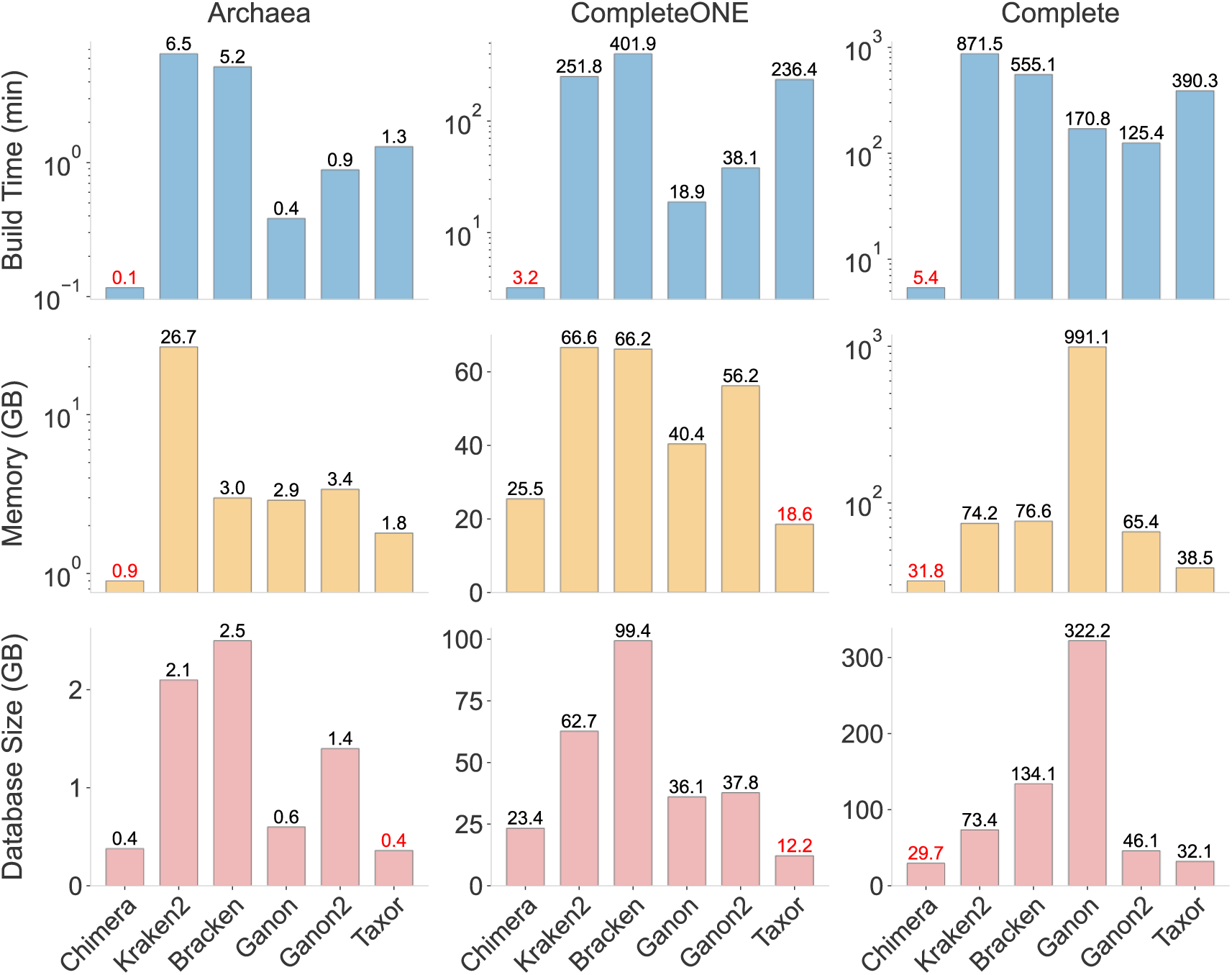
Performance benchmarking and detailed information for different taxonomic classifiers and reference databases. (A) Database construction comparison among six classification tools (Chimera, Kraken2, Bracken, Ganon, Ganon2, and Taxor) using three distinct datasets (Archaea, CompleteONE, and Complete). The metrics evaluated are database build time (top panels, blue), peak memory usage (middle panels, yellow), and final database size (bottom panels, red). Optimal (lowest) values in each category are highlighted in red text.

Chimera exhibited superior performance across all datasets, most notably being the only tool capable of successfully constructing the full RefSeq database within a 1 TB memory constraint, completing the task in approximately two hours; all other tools failed due to memory overflow. For the CompleteONE dataset, Chimera achieved a 78-fold faster construction time compared to Kraken2 (98.7% reduction) and a 74-fold improvement compared to Taxor (98.6% reduction). On the larger Complete dataset, Chimera’s construction time was 162-fold faster than Kraken2 (99.4% reduction) and 74-fold faster than Taxor (98.6% reduction). Furthermore, Chimera required only 31.8 GB of memory and produced a 29.7 GB database for the Complete dataset—markedly lower than Kraken2’s memory usage of 74.2 GB and a database size of 73.4 GB. This significant reduction in memory usage makes high-quality database construction feasible even on personal computers.

Overall, Chimera consistently demonstrates unparalleled efficiency and scalability, standing as the only tool capable of constructing the entire RefSeq database within a 1 TB memory constraint. Most notably, Chimera achieves what no other classifier can—updating the widely used Complete database in an astonishing five minutes. This revolutionary speed redefines the feasibility of daily database updates, eliminating the hours or even days required by existing tools. By combining unmatched computational efficiency with minimal memory demands, Chimera shatters traditional hardware limitations, democratizing large-scale metagenomic analysis across all research environments. As a result, Chimera is not just an incremental improvement but a paradigm shift in metagenomic classification, offering unprecedented support for high-resolution microbial community analysis and future microbiome research.

### 2.3 Classification Performance on Constructed Databases

This section evaluates Chimera’s classification performance using databases constructed in previous experiments, comparing it against widely-used tools such as Kraken2, Bracken, Ganon, Ganon2, and Taxor. Classification assessments were conducted using both Complete and CompleteONE databases to ensure consistency and comparability.

Four real and simulated datasets from the CAMI II project were employed, supplemented by an additional simulated dataset to enrich the evaluation (Supplemental Table S3) [27,28]. These datasets encompass diverse sequencing strategies, including long-read (average ∼3,000 bp) and short-read (2 × 150 bp) sequences, designed to simulate complex microbial ecosystems such as marine and mouse gut microbiomes. The simulated datasets included technical sequencing errors and random insert-size variations to evaluate robustness and adaptability of classification algorithms.

Experiments were performed under the same hardware conditions as database construction, with all tools executed using default or recommended settings, a uniform classification threshold of 70%, and a fixed configuration of 32 computational threads to ensure a fair comparison. Performance metrics included accuracy, precision, recall, F1-score, and L1 distance (detailed calculation methods are provided in the supplementary materials). Notably, L1 distance quantifies the discrepancy between predicted and true abundances, with lower values indicating higher precision and reliability in ecological community profiling.

Results indicate that Chimera consistently delivered outstanding performance across most metrics (Table 1, Supplemental Figure S1). Chimera achieved the highest accuracy and F1 scores across nearly all datasets, except for the CAMI Marine long-read dataset using the CompleteONE database, where Ganon slightly outperformed Chimera. Although Kraken2 demonstrated superior precision, it significantly lagged in accuracy, recall, and F1-score. Ganon achieved the highest recall in specific datasets, yet Chimera maintained stable recall performance overall. Importantly, Chimera consistently exhibited the lowest L1 distance across all datasets, underscoring its precision and reliability in abundance estimation.

**Table 1.** Taxonomic Classification Performance on Benchmark Datasets.

| Software | Dataset Name | CompleteONE |  |  |  | Complete |  |  |  |
| --- | --- | --- | --- | --- | --- | --- | --- | --- | --- |
|  |  | Accuracy | Precision | Recall | F1 Score | Accuracy | Precision | Recall | F1 Score |
| **Chimera** | CAMI II Marine (long read) | 0.50 | 0.87 | 0.54 | 0.66 | 0.60 | 0.91 | 0.64 | 0.75 |
| **Kraken2** |  | 0.38 | 0.98 | 0.39 | 0.55 | 0.44 | 0.98 | 0.44 | 0.61 |
| **Ganon** |  | 0.50 | 0.87 | 0.54 | 0.67 | 0.40 | 0.55 | 0.60 | 0.57 |
| **Ganon2** |  | 0.49 | 0.85 | 0.54 | 0.66 | 0.59 | 0.89 | 0.63 | 0.74 |
| **Taxor** |  | 0.45 | 0.88 | 0.48 | 0.62 | 0.57 | 0.92 | 0.60 | 0.73 |
| **Chimera** | CAMI II Marine (short read) | 0.52 | 0.86 | 0.57 | 0.68 | 0.61 | 0.90 | 0.66 | 0.76 |
| **Kraken2** |  | 0.39 | 0.97 | 0.39 | 0.56 | 0.44 | 0.98 | 0.45 | 0.61 |
| **Ganon** |  | 0.51 | 0.85 | 0.57 | 0.68 | 0.48 | 0.65 | 0.64 | 0.65 |
| **Ganon2** |  | 0.50 | 0.83 | 0.56 | 0.67 | 0.59 | 0.86 | 0.65 | 0.74 |
| **Taxor** |  | 0.47 | 0.87 | 0.51 | 0.64 | 0.59 | 0.90 | 0.63 | 0.74 |
| **Chimera** | CAMI II Toy Mouse Gut (long read) | 0.28 | 0.74 | 0.31 | 0.44 | 0.35 | 0.74 | 0.39 | 0.52 |
| **Kraken2** |  | 0.20 | 0.84 | 0.21 | 0.33 | 0.25 | 0.85 | 0.26 | 0.40 |
| **Ganon** |  | 0.28 | 0.73 | 0.31 | 0.44 | 0.28 | 0.43 | 0.43 | 0.43 |
| **Ganon2** |  | 0.27 | 0.71 | 0.30 | 0.43 | 0.34 | 0.75 | 0.39 | 0.51 |
| **Taxor** |  | 0.17 | 0.73 | 0.18 | 0.29 | 0.28 | 0.76 | 0.31 | 0.44 |
| **Chimera** | CAMI II Toy Mouse Gut (short read) | 0.28 | 0.73 | 0.31 | 0.43 | 0.35 | 0.75 | 0.39 | 0.51 |
| **Kraken2** |  | 0.20 | 0.83 | 0.21 | 0.33 | 0.25 | 0.85 | 0.26 | 0.40 |
| **Ganon** |  | 0.27 | 0.72 | 0.30 | 0.43 | 0.29 | 0.48 | 0.43 | 0.45 |
| **Ganon2** |  | 0.26 | 0.70 | 0.30 | 0.42 | 0.34 | 0.74 | 0.38 | 0.50 |
| **Taxor** |  | 0.18 | 0.72 | 0.19 | 0.30 | 0.30 | 0.77 | 0.32 | 0.46 |
| **Chimera** | Simulated Dataset | 0.58 | 0.67 | 0.81 | 0.73 | 0.88 | 0.90 | 0.97 | 0.94 |
| **Kraken2** |  | 0.24 | 0.99 | 0.24 | 0.39 | 0.30 | 0.99 | 0.30 | 0.46 |
| **Ganon** |  | 0.56 | 0.65 | 0.80 | 0.72 | 0.59 | 0.60 | 0.99 | 0.74 |
| **Ganon2** |  | 0.54 | 0.63 | 0.80 | 0.70 | 0.71 | 0.72 | 0.98 | 0.83 |
| **Taxor** |  | 0.27 | 0.43 | 0.42 | 0.42 | 0.79 | 0.87 | 0.89 | 0.88 |

Results showed that Chimera exhibited moderate runtimes, outperforming Ganon and Kraken2 but slightly behind Ganon2 and Taxor, while demonstrating significantly better memory efficiency, slightly lower than Taxor and markedly better than the other tools (Supplemental Figure S2). Except for Kraken2, memory consumption was primarily driven by database size rather than the volume of classified sequences. Taxor’s advantage in memory usage was mainly attributed to its smaller CompleteONE database.

In summary, Chimera achieves exceptional classification performance while maintaining outstanding database construction efficiency, surpassing or matching the best-performing tools across all key metrics. Notably, Chimera combines superior accuracy with minimal computational resource demands, rendering it highly suitable for large-scale data processing and environments with constrained computational resources. These attributes establish Chimera as a leading and forward-looking metagenomic classification solution, providing unparalleled technical support for high-throughput microbiome research.

### 2.4 Effects of FMC on Classification Performance

In this study, we introduced FMC, a novel database balancing strategy that significantly improves metagenomic classification by systematically limiting the number of minimizer hashes per species. FMC effectively mitigates taxonomic overrepresentation, a prevalent issue wherein abundant species disproportionately dominate hash representation, masking signals from less abundant taxa and consequently diminishing overall classification accuracy [16,18]. To rigorously evaluate FMC’s impact, we constructed a series of databases with varying maximum hash limits using the Complete dataset and assessed classification performance using the CAMI II Marine long-read dataset (Figure 3). Results revealed substantial sensitivity of classification performance to hash limit parameters (Figure 3B). Classification accuracy and F1 scores improved sharply at a hash limit of 2×10^5^ and peaked at 5×10^5^, beyond which performance declined, indicating that excessive hash inclusion introduces noise and redundancy. Databases constructed without hash constraints exhibited significantly inferior classification outcomes alongside larger database sizes (Figure 3A), highlighting the critical importance of controlled hash allocation. Consequently, we selected a 2×10^5^ hash limit as Chimera’s default parameter for the Complete dataset, achieving near-optimal classification performance while maintaining minimal memory and storage requirements. Under the 2×10^5^ hash limit, the maximum memory usage per species can be estimated as 3.81 MB. However, in practical applications, due to the filtering and deduplication of FMC, most species complete genome do not reach this maximum limit; consequently, the average actual memory usage per species is approximately 1.21 MB. Under these conditions, a system with 1 TB of memory is estimated to accommodate over 800,000 species, demonstrating the scalability of FMC in handling the growing complexity of metagenomic datasets.On larger and more complex datasets like RefSeq, increased hash limits further enhanced classification effectiveness; for example, the 4×10^5^ hash limit substantially outperformed the 2×10^5^ limit (Figure 3C). Nonetheless, even the 2×10^5^ setting on the RefSeq dataset yielded classification outcomes comparable to those of the Complete dataset, with slightly higher accuracy and F1 scores. These findings underscore that although higher hash limits may benefit extremely comprehensive datasets by preserving more critical information, lower hash limits remain highly effective, significantly reducing resource consumption.

**Figure 3.**
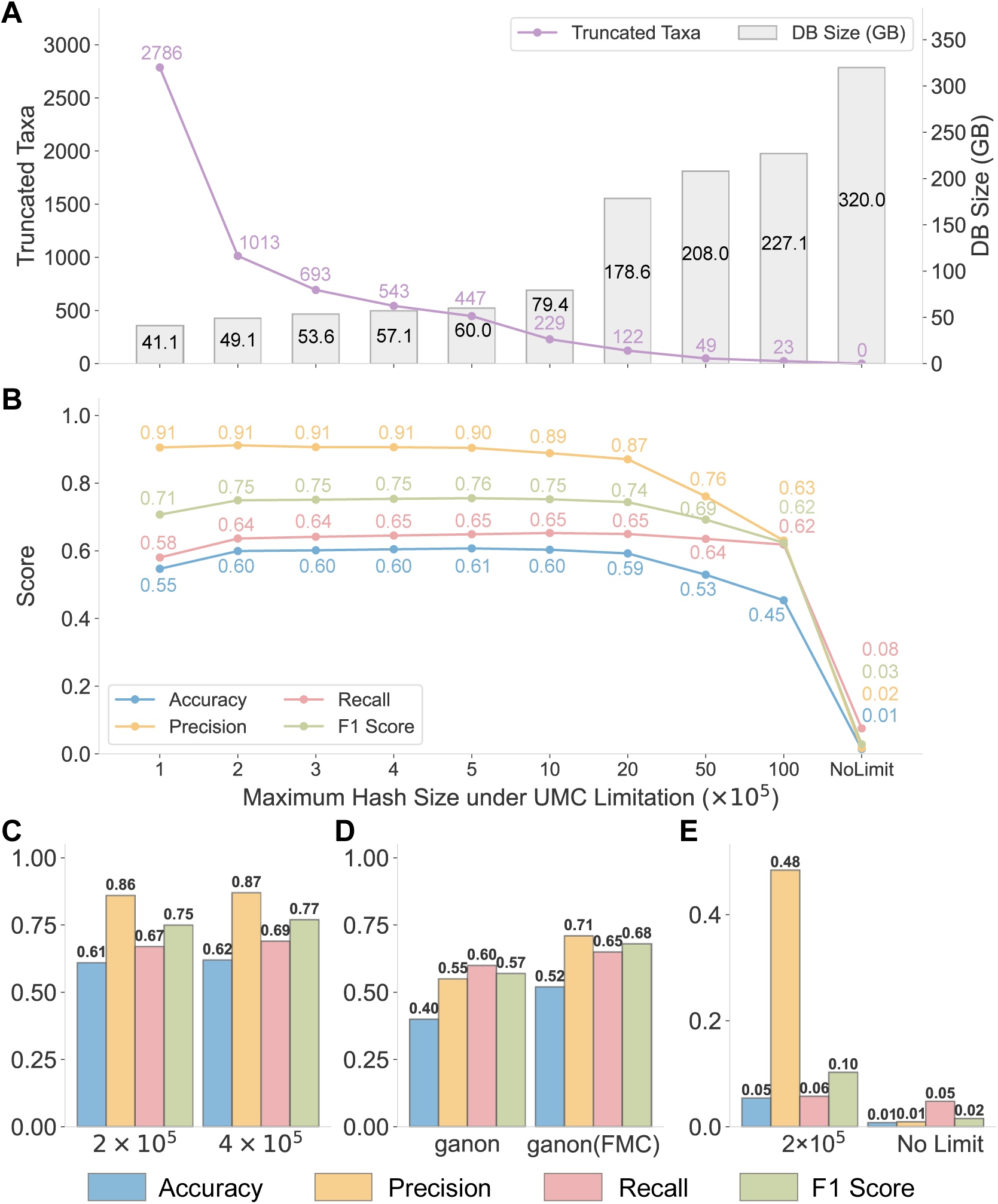
Impact of FMC on metagenomic classification performance using the CAMI II Marine long-read dataset. (A) Number of truncated taxa (purple line) and associated database size (grey bars) across varying maximum hash sizes (1×10^5^ to 100×10^5^) and in an unconstrained (“No Limit”) scenario. (B) Classification performance (accuracy, precision, recall, and F1 score) across different hash size constraints under the complete dataset, illustrating the balance between accuracy and database efficiency. (C) Performance metrics for selected hash sizes (2×10⁵ and 4×10⁵) using the RefSeq dataset. (D) Performance comparison of ganon classifier with and without FMC (ganon vs ganon(FMC)) on the complete dataset. (E) Classification metrics for 54 randomly selected truncated taxa, comparing fixed hash size (2×10⁵) and unconstrained scenarios under the complete dataset.

To further validate the versatility of FMC, we integrated it into the ganon classifier by modifying its source code and incorporating FMC into its database construction pipeline (Supplementary Materials). This experiment demonstrated substantial performance gains across all evaluated metrics (Figure 3D), confirming FMC’s broad applicability beyond Chimera and highlighting its potential to enhance the classification accuracy and resource efficiency of various metagenomic tools.

Given that the 2×10^5^ hash limit led to the truncation of 1013 taxa (Figure 3A), we designed a targeted validation experiment to assess whether truncation negatively affected classification accuracy. We randomly selected 54 truncated taxa, downloading approximately ten sequences per taxon from NCBI, and generated an independent test dataset. Remarkably, classification performance for these truncated taxa under the 2×10^5^ FMC condition remained superior to the scenario without FMC. Critically, the detrimental impact of taxonomic overrepresentation outweighed any potential negative effects of hash truncation. By balancing hash distribution, FMC substantially improved the detectability of low-abundance taxa signals, enhancing overall classification robustness and stability (Figure 3E). Thus, rather than weakening classification capabilities, targeted hash truncation through FMC effectively counteracted biases from taxonomic overrepresentation.

In conclusion, FMC is a transformative strategy that not only optimizes database size, memory efficiency, and classification performance but also provides a universal solution to the long-standing issue of taxonomic overrepresentation. By systematically balancing species representation, FMC eliminates biases inherent in traditional metagenomic classification, setting a new standard for database construction. Moreover, the powerful synergy of FMC and IMCF delivers a solution to the challenge posed by the exponential growth of species in genomic databases. This framework is not just sufficient for current metagenomic datasets but is fully equipped to scale with the future explosion of genomic data. As large-scale sequencing efforts continue to expand, FMC and IMCF together establish a robust and forward-looking foundation for the next generation of metagenomic research.

## 3. Discussion

Our results demonstrate Chimera’s exceptional efficiency in database construction and robust performance in metagenomic classification, positioning it as an essential tool for contemporary microbiome research. A remarkable advantage of Chimera lies in its extraordinary speed and low memory footprint during database construction. Notably, Chimera can construct a complete RefSeq genome database in approximately five minutes using less than 32 GB of memory, and complete the entire RefSeq database construction within approximately two hours. This unprecedented efficiency significantly reduces dependence on advanced computational infrastructure, enabling high-quality metagenomic analyses to be performed even on standard laboratory equipment or personal computers, substantially broadening the accessibility and applicability of metagenomics research [5,27]. The ability to rapidly update databases makes Chimera particularly valuable in research environments that require frequent reference updates, such as pathogen surveillance, clinical diagnostics, and environmental monitoring, ensuring that classification tools always operate with the most up-to-date genomic data [2,29].

Additionally, Chimera’s database scalability is predictable, and its maximum storage requirements increase linearly with the number of species included. Under default parameters, Chimera allocates at most 3.81 MB per species, but in actual use, the average memory requirement is approximately 1.21 MB per species. As a result, a 1 TB memory system is expected to accommodate databases containing over 800,000 species. By comparison, as of March 10, 2025, the RefSeq database comprises data from only 164,117 species, suggesting that Chimera’s architecture can comfortably accommodate database expansions for at least the next decade. This robust scalability not only addresses current computational bottlenecks but also establishes a solid foundation for handling increasingly complex microbial datasets in the future.

Chimera’s superior performance is primarily driven by two core innovations: IMCF and FMC. The IMCF, an advanced Bloom-filter variant, significantly enhances classification accuracy through highly efficient minimizer indexing, remarkably low false-positive rates, and SIMD-accelerated sequence queries, leading to substantial improvements in both classification speed and precision. The FMC strategy effectively mitigates biases arising from taxonomic overrepresentation by strictly limiting the number of hashes allocated per species, thereby substantially reducing memory consumption and accelerating database construction. Furthermore, the general applicability of FMC was demonstrated by successfully integrating it into another classification tool, ganon, resulting in significant improvements across multiple classification metrics [12]. Thus, FMC not only optimizes Chimera’s performance but can broadly enhance other hash-based metagenomic classifiers.

The synergy of IMCF and FMC strategies allows Chimera to surpass mainstream classification tools, such as Kraken2 and Taxor, achieving consistently higher accuracy and F1 scores, and significantly reducing errors in abundance estimation (L1 distance). This capability makes Chimera particularly suited for investigating microbial diversity and ecological functions, especially in reliably identifying rare or low-abundance microorganisms commonly overlooked by traditional classification approaches.

Despite its substantial advancements, Chimera presents areas that warrant further optimization. Currently, database construction requires manual tuning of the load factor, where inappropriate settings could either lead to construction failures or unnecessarily large databases. Moreover, the current minimizer selection mechanism within FMC is relatively simplistic, limiting the potential optimization of k-mer informativeness and thus hindering further improvements in classification performance. Addressing these limitations should be a key focus of future research, including the development of automated algorithms for optimal load-factor determination to streamline database construction. Additionally, integrating statistical or machine learning approaches for improved minimizer selection could significantly enhance classification accuracy. Enhancements in data insertion methods and spatial efficiency would further increase Chimera’s scalability to meet the demands of increasingly large and dynamically evolving databases. Furthermore, our research group has already established a robust foundation in sequence alignment, providing a strong platform to potentially incorporate pangenome graphs into Chimera in the future [30–32]. This integration would enable precise strain-level classification, further elevating Chimera’s analytical resolution [33,34].

Collectively, these technological advancements will substantially expand Chimera’s application potential, enabling it to adapt effectively not only to diverse and resource-limited research scenarios but also to future large-scale, complex metagenomic data analyses. Chimera’s remarkable computational efficiency and scalability position it as a foundational tool for next-generation metagenomic research, offering researchers globally a sustainable, efficient framework for database construction and microbial classification. Continued refinement and innovation will likely establish Chimera as a standard analytical tool in metagenomics, propelling high-resolution microbiome research and providing robust technological support for deeper explorations into microbial dynamics, evolutionary patterns, and ecosystem functionality.

## 4. Conclusions

We introduce Chimera, a highly efficient and precise metagenomic classification tool designed to address critical computational and database construction challenges in microbial research. Leveraging two key innovations—the IMCF and the FMC strategy—Chimera achieves exceptional classification accuracy, rapid query speeds, and significant memory reduction. It can construct the RefSeq complete genome database, within approximately five minutes using less than 32 GB of memory, while also ensuring scalable storage (up to 800,000 species per 1 TB memory). Experimental results highlight Chimera’s superior accuracy, especially in identifying low-abundance taxa, alongside the broad applicability of FMC in optimizing other classification tools. Overall, Chimera provides a robust, scalable, and accessible framework for next-generation metagenomic research, enabling deeper exploration into microbial diversity and ecological interactions.

## 5. Methods

### 5.1 Interleaved Cuckoo Filter

The Interleaved Bloom Filter (IBF) is a widely used data structure in large-scale metagenomic classification due to its ability to perform simultaneous queries across multiple Bloom filters with high efficiency, enabled by its interleaved encoding scheme [12,24,35]. This makes IBF particularly suitable for high-throughput applications. However, IBF’s practical utility is limited by several inherent drawbacks: its construction process is computationally expensive, its false positive rate remains relatively high, and its interleaved structure imposes a rigid uniformity constraint, requiring all filters to have the same size. This constraint often results in significant memory overhead in imbalanced datasets, as smaller taxa must conform to the size of the largest taxa.

The Interleaved Cuckoo Filter (ICF) builds upon the principles of IBF, addressing these challenges by significantly improving construction speed and query accuracy[23]. By leveraging the adaptability of Cuckoo hashing, ICF reduces computational overhead during filter construction, while its splitting mechanism for large taxa mitigates the inefficiencies caused by dataset imbalance. Integrating multiple Cuckoo Filters into a single interleaved bit-vector array, ICF maintains a compact design that scales effectively for large-scale, high-throughput genomic datasets. Figure 4 provides an overview of its structure.

**Figure 4.**
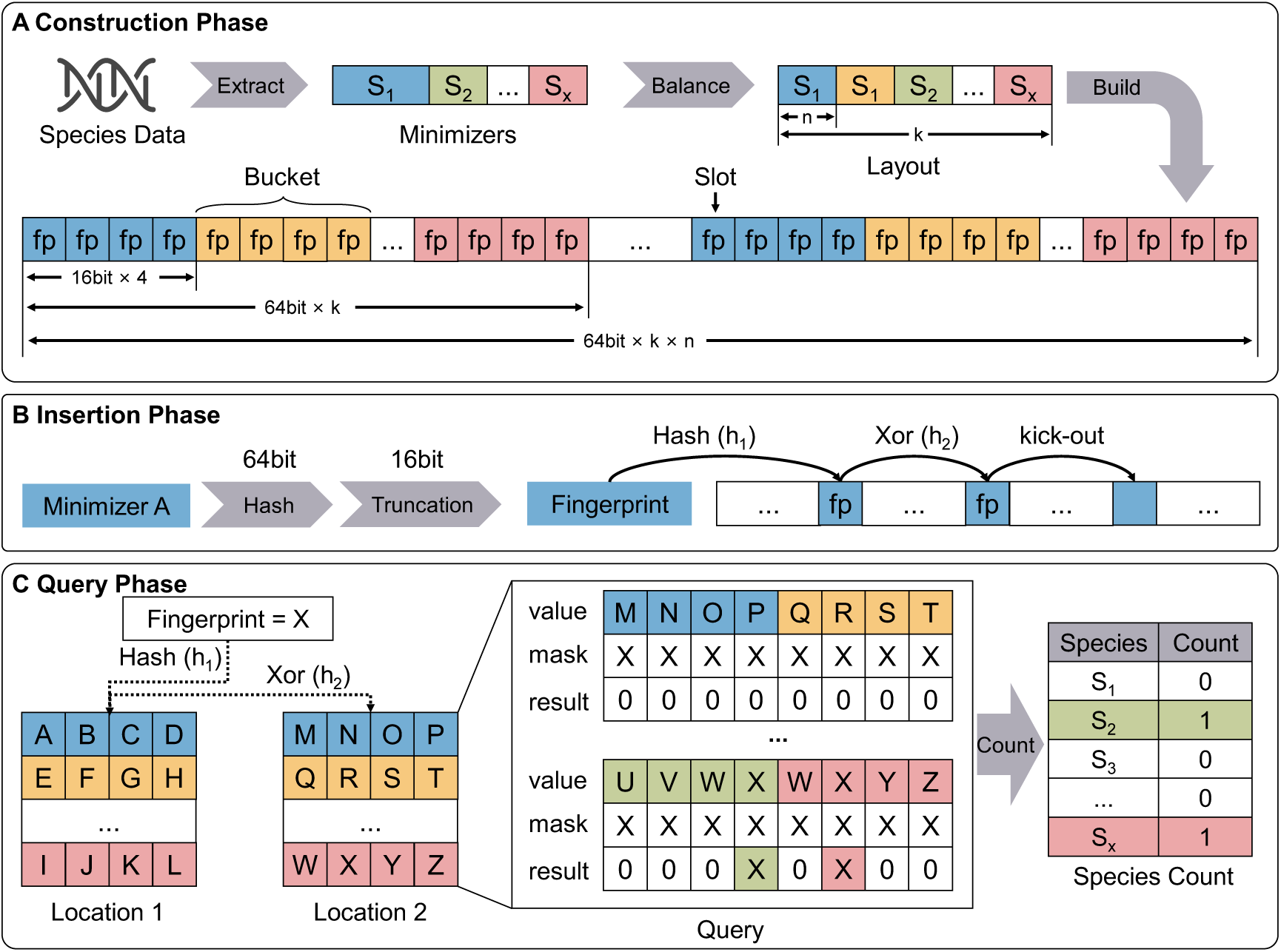
Construction and Query Workflow of the ICF. (A) Construction Phase: Extracts minimizers from input data, performs load balancing, and builds the ICF structure. (B) Insertion Phase: Computes the hash value of a minimizer, truncates it to obtain a fingerprint, and inserts it into the ICF structure. If both candidate positions computed by the hash functions are occupied, a kick-out operation is performed, evicting one fingerprint and relocating it to an alternate position to optimize storage efficiency. (C) Query Phase: Utilizes a mask-based approach to simultaneously query eight slots, improving fingerprint matching efficiency and reducing query latency.

As shown in Figure 4A, the construction of ICF employs a binary search algorithm to determine the optimal number and size of Cuckoo Filters, ensuring balanced load distribution while minimizing memory overhead. Specifically, let *M* denote the total number of minimizers to be inserted, *α* represent the predefined load factor controlling the fill ratio of each filter, *k* denote the number of Cuckoo Filters, and *n* denote the capacity of a single filter. ICF aims to satisfy the condition:

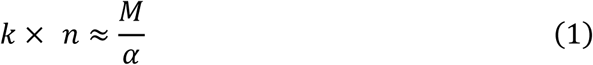

ensuring that the total storage capacity *k* × *n* approximates the dataset size scaled by the inverse of the load factor, thereby minimizing unnecessary overhead.

The binary search begins with a lower bound of zero and an upper bound set to twice the number of minimizers in the largest taxon, iteratively refining the range until convergence to the optimal configuration satisfying Equation 1. During this process, oversized filters are considered for splitting dynamically, ensuring that the number of filters is determined by memory efficiency constraints rather than being solely dictated by the number of taxa.

The choice of the load factor *α* is critical: a higher *α* improves memory efficiency but may increase insertion conflicts, potentially leading to degraded performance or insertion failures. Conversely, a lower *α* reduces conflicts but results in excessive memory overhead.

After the construction phase determines the optimal number and size of Cuckoo Filters, ICF proceeds with the insertion phase, where each element is mapped to two candidate bucket positions using two independent hash functions. The primary hash function *h*_1_ is defined as:

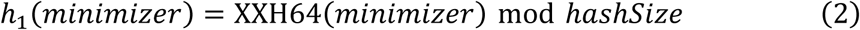

where the minimizer is a 64-bit unsigned integer encoding a k-mer representative, and *hashSize* represents the number of buckets in a single Cuckoo Filter. XXH64 is a variant of the xxHash (https://github.com/Cyan4973/xxHash) family, known for its high-speed and high-quality hashing performance. The secondary hash function *h*_2_ is computed as:

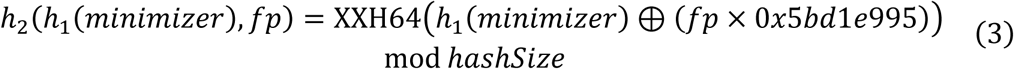

where *fp* is a 16-bit fingerprint derived from the minimizer. The multiplication by the constant 0*x*5*bd*1*e*995 enhances randomness and improves hash dispersion, reducing insertion conflicts and potential clustering effects.

As illustrated in Figure 4B, if both candidate buckets are occupied, the insertion procedure initiates a “kick-out” operation, relocating an existing element to its alternate bucket. This process repeats iteratively until an empty slot is found or a rehashing threshold is reached. Leveraging the efficiency of Cuckoo hashing, ICF maintains a low storage overhead while ensuring robust insertion performance.

The querying process in ICF follows a similar pattern as insertion, leveraging the same hash functions and interleaved data structure for efficient lookups. Given a query minimizer and its fingerprint the algorithm computes its two candidate bucket positions using the previously defined hash functions (Equations 2 and 3), then checks whether the corresponding interleaved buckets contain a matching fingerprint. To accelerate lookups, ICF’s interleaved design allows multiple Cuckoo Filters to be stored within a single bit-vector, enabling SIMD-based acceleration via AVX2 instructions for parallel 16-bit fingerprint comparison as shown in Figure 4C. Additionally, the movemask instruction extracts results in a single operation, reducing branch misprediction overhead and significantly improving query performance. This vectorized lookup mechanism effectively reduces query latency compared to traditional sequential lookups.

Despite the improvements in storage efficiency and query accuracy, ICF faces scalability challenges when applied to large-scale genomic datasets. As the dataset size expands, the number of required Cuckoo Filters increases, leading to higher lookup latency. Additionally, the use of 16-bit fingerprints, while improving accuracy, introduces additional memory overhead compared to Bloom filters, potentially affecting overall efficiency. Addressing these limitations requires further optimization, particularly in accelerating classification and managing database scalability.

### 5.2 Interleaved Merged Cuckoo Filter

To overcome the bottlenecks of the ICF in terms of classification speed and construction efficiency, we propose an improved data structure: the IMCF. This method introduces multiple optimizations for large-scale data processing in metagenomic analysis, significantly enhancing storage efficiency and query performance. The core idea of IMCF is to utilize the first 4 bits of the 16-bit fingerprint for storing species index information while retaining the last 12 bits for the minimizer hash fingerprint (as illustrated in Figure 5A). Let *I* denote the species index (0 ≤ *I* < 16), and *h*(⋅) represent the hash function that generates the 12-bit fingerprint. Then, each inserted fingerprint can be expressed as:

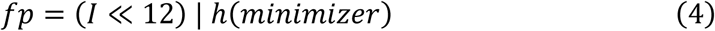

**Figure 5.**
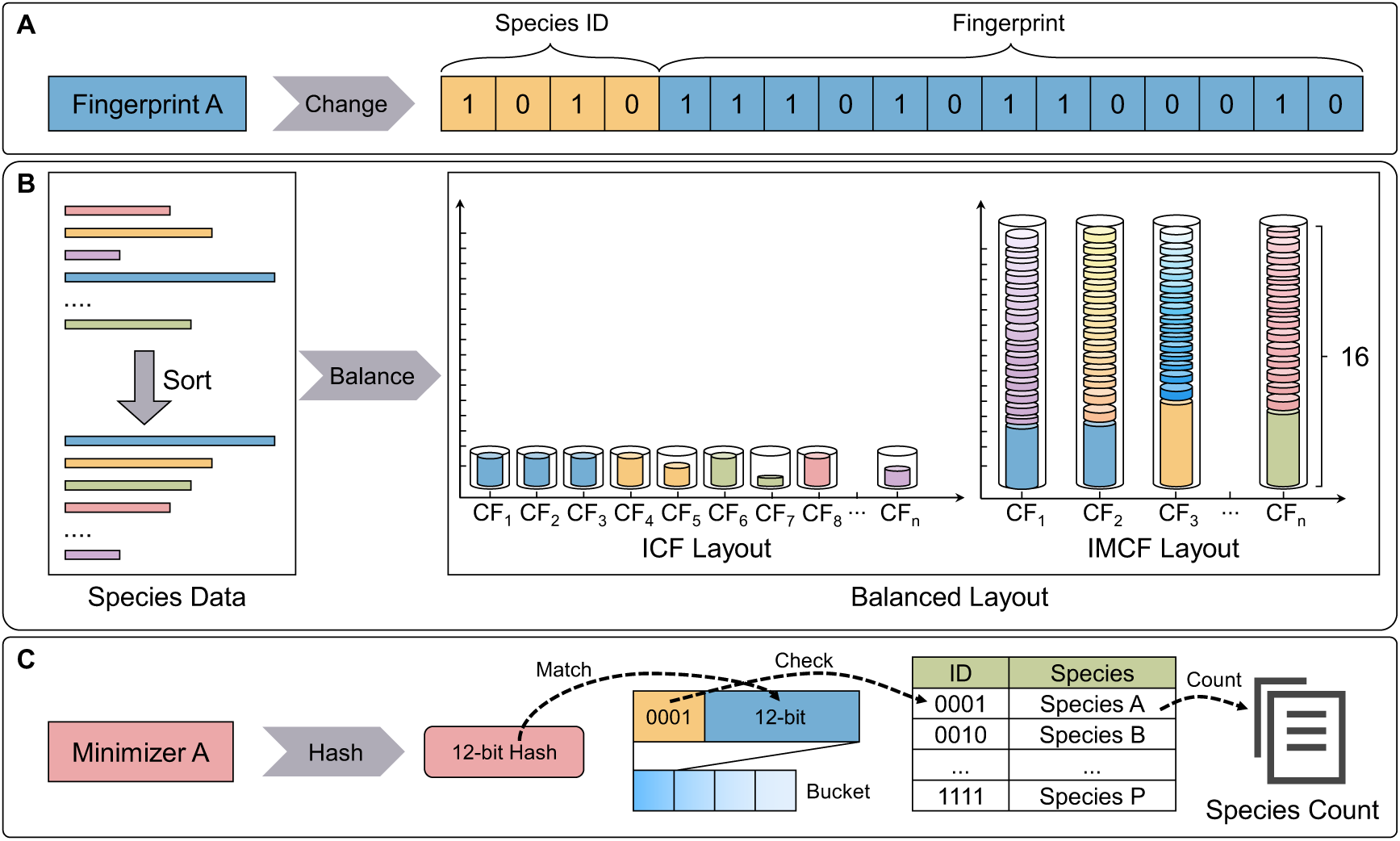
Design, Construction, and Query Process of the IMCF. (A) To optimize storage and querying, the original 1-bit fingerprint from the ICF is split into a 4-bit species ID and a 12-bit fingerprint. The 4-bit species ID allows the IMCF to handle up to 1 species within a single filter, while the 12-bit fingerprint retains the specificity required for accurate identification. (B) During the construction process, species data are first sorted in descending order by size. Then, a greedy algorithm is used to distribute the data across different IMCF cuckoo filters, ensuring balanced data distribution and improved query efficiency. The figure compares the layout of the cuckoo filters in ICF and IMCF. (C) During querying, a 4-bit minimizer is first hashed into a 12-bit fingerprint. The IMCF then begins searching for matches in the interleaved slots. If the 12-bit portion of a slot matches the query, the 4-bit species ID stored in the same slot is checked to verify the corresponding species.

where “≪” denotes the left shift operation, and “|” represents the bitwise OR operation. Through this design, a single query can simultaneously match up to 16 species by first verifying the last 12 bits of the fingerprint; if a match is found, the first 4-bit species index is subsequently checked (as illustrated in Figure 5C). This approach leads to more than a 16-fold increase in query efficiency, while the integration of the species index within the fingerprint also simplifies the insertion process, accelerating filter construction.

During the storage construction phase, IMCF ensures efficient load balancing by maintaining uniform filter sizes while concurrently storing multiple species within each Cuckoo Filter (as illustrated in Figure 5B). Specifically, the number of minimizer hashes for all species, denoted as {*M*_1_, *M*_2_,…, *M*_n_}, is first computed to determine the median value *Med*. A threshold for splitting species into smaller blocks is then derived as:

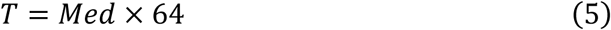

If the minimizer count for a species *M*_i_ exceeds *T*, the species data is split into smaller hash blocks for distribution among different filters. A greedy strategy is employed in this process: the largest minimizer hash block is always assigned to the filter with the lowest current storage load, ensuring balanced storage distribution across filters. This design enables IMCF to maintain efficient memory utilization while mitigating excessive insertion failures and storage imbalance, even in large-scale and highly complex datasets.

It is noteworthy that since each filter simultaneously stores up to 16 species while using the same 12-bit fingerprint space for position calculations, the probability of hash collisions increases, potentially leading to higher insertion failure rates under extreme loads. To address this issue, IMCF adopts a lower load factor, reserving additional free space within the filter to alleviate collisions and reduce insertion failures. Although decreasing the load factor slightly increases the storage overhead per filter, IMCF still achieves a significantly higher compression ratio than conventional ICF while leveraging parallelism to achieve a remarkable improvement in query performance. Therefore, IMCF effectively balances the trade-offs between hash collision risks and storage efficiency, offering a highly efficient solution for large-scale metagenomic data retrieval with enhanced classification accuracy and construction speed.

### 5.3 Database Construction

The Chimera database construction pipeline is designed to maximize classification efficiency and accuracy while minimizing resource consumption. At its core is the FairMin-Cap (FMC) strategy, which optimizes minimizer selection, controls database size, and ensures balanced species representation, thus significantly enhancing downstream classification performance.

The pipeline begins with data retrieval using genome_updater (https://github.com/pirovc/genome_updater), directly acquiring reference genomes and associated sequences from NCBI without additional preprocessing.

Minimizers, compact representations derived from k-mers, are then extracted from the raw datasets, effectively reducing memory usage and improving query efficiency.

The FMC strategy employs adaptive file-size thresholds to filter low-frequency minimizers. For compressed files, threshold assignment is based on the estimated decompressed file size (s), as follows:

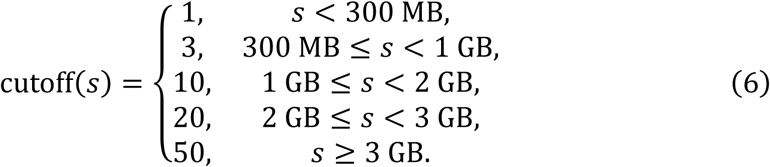

This method retains essential classification information while significantly reducing noise. Subsequently, FMC removes duplicate minimizers to further reduce redundancy and sets a default upper limit of two million minimizers per species. By limiting the representation of highly abundant species, FMC effectively controls database size, maintains taxonomic balance, significantly enhances classification accuracy and query speed, and reduces overall memory usage.

After minimizer-related processing, Chimera configures the IMCF indexing structure using the greedy optimization algorithm detailed in Section 5.2. This structure stores minimizer data alongside corresponding taxonomic labels, enabling rapid and accurate downstream classification analysis.

### 5.4 Sequence Classification

Chimera’s sequence classification pipeline is engineered to handle the complexity and scalability demands of metagenomic data analysis. The workflow integrates high-efficiency minimizer matching, stringent filtering steps, and adaptable classification algorithms to optimize accuracy and computational performance.

The process initiates by extracting minimizers from input sequences and rapidly matching them against a preconstructed database stored in an IMCF. This structure supports fast query operations and ensures effective utilization of computational resources. Once the minimizer matching is completed, the system refines the classification results through a four-step filtering procedure. First, matches below a predefined threshold are discarded to eliminate low-confidence signals. Next, matches contributing less than 80% of the maximum match count are excluded to minimize noise and enhance signal-to-noise ratios. Subsequently, a list of reference genomes containing at least one uniquely matched read segment is constructed, and reads assigned to genomes lacking such matches are removed.

For reference genomes with fewer than 5% uniquely matched reads that share 95% of matches with another genome, all matches are reassigned to the dominant genome to further improve classification precision[36].

Chimera employs three classification algorithms, each tailored for distinct analytical requirements, with one algorithm selectable per analysis. The default method is VEM (Variational EM), which extends Expectation-Maximization with Bayesian inference to refine abundance estimates and improve classification reliability. LCA (Lowest Common Ancestor) assigns sequences to the lowest taxonomic rank shared by matching references, favoring conservative assignments ideal for biodiversity profiling. EM (Expectation-Maximization) iteratively estimates sequence abundances, making it suitable for resolving complex abundance distributions.

During the final optimization stage, Chimera evaluates the outputs of the selected algorithm, correcting biases in abundance estimates and computing confidence intervals to assess classification uncertainty. Results are provided for individual sequences and can be aggregated to produce global abundance estimates. Chimera also facilitates visual exploration of classification outputs through Krona, enabling hierarchical visualization of taxonomic composition and abundance patterns [21]. Outputs are formatted in standardized file types to ensure compatibility with downstream analytical tools.

## Data availability

All data and code used in this study are publicly available. The source code for Chimera can be accessed at https://github.com/malabz/Chimera, while benchmarking scripts and related information are provided at https://github.com/malabz/ChimeraBenchmark. The RefSeq dataset used for database construction was retrieved using genome_updater (https://github.com/pirovc/genome_updater), which can also be invoked directly within Chimera for dataset acquisition.

For metagenomic classification, we utilized multiple datasets, including the CAMI II Toy Mouse Gut dataset (https://frl.publisso.de/data/frl:6421672/), the CAMI II Marine dataset (https://frl.publisso.de/data/frl:6425521/marine/), and a simulated dataset available at https://doi.org/10.5281/zenodo.10666087.

All experimental results supporting this study are deposited at https://doi.org/10.5281/zenodo.15081818.

## Acknowledgment

This work was supported by the National Natural Science Foundation of China (Grant No. 62373080) and Zhongguancun Academy, under the research projects Project No. 20240310 and Project No. 20240101. We sincerely appreciate their support in funding and resources.

## Author contributions

Qinzhong Tian and Pinglu Zhang contributed equally to this work. Qinzhong Tian led the project, designed the methodology, and supervised the research. Pinglu Zhang contributed to algorithm development, data analysis, and experimental validation. Yanming Wei assisted with software implementation and performance benchmarking. Quan Zou provided theoretical guidance and contributed to manuscript revision. Yansu Wang and Ximei Luo jointly supervised the study, provided critical insights, and were responsible for securing funding. All authors reviewed and approved the final manuscript.

## Notes

https://doi.org/10.5281/zenodo.15081818

https://github.com/malabz/Chimera

https://github.com/malabz/ChimeraBenchmark

### Competing Interest Statement

The authors have declared no competing interest.

