## Supplemental Information for "Chimera: Ultrafast and Memory-efficient Database Construction for High-Accuracy Taxonomic Classification in the Age of Expanding Genomic Data"

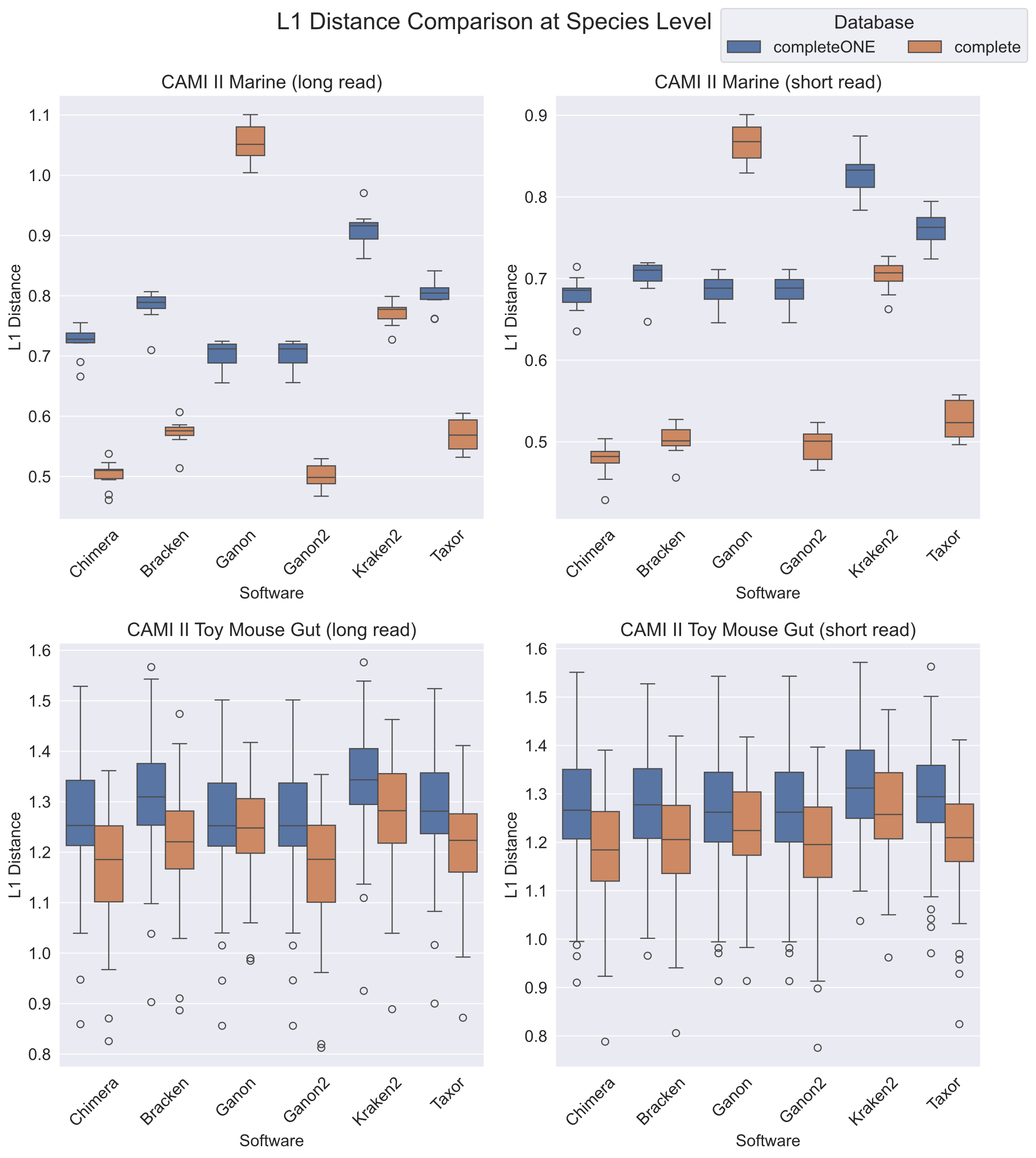

Figure S1 L1 Distance Comparison Across Tools at Species Level

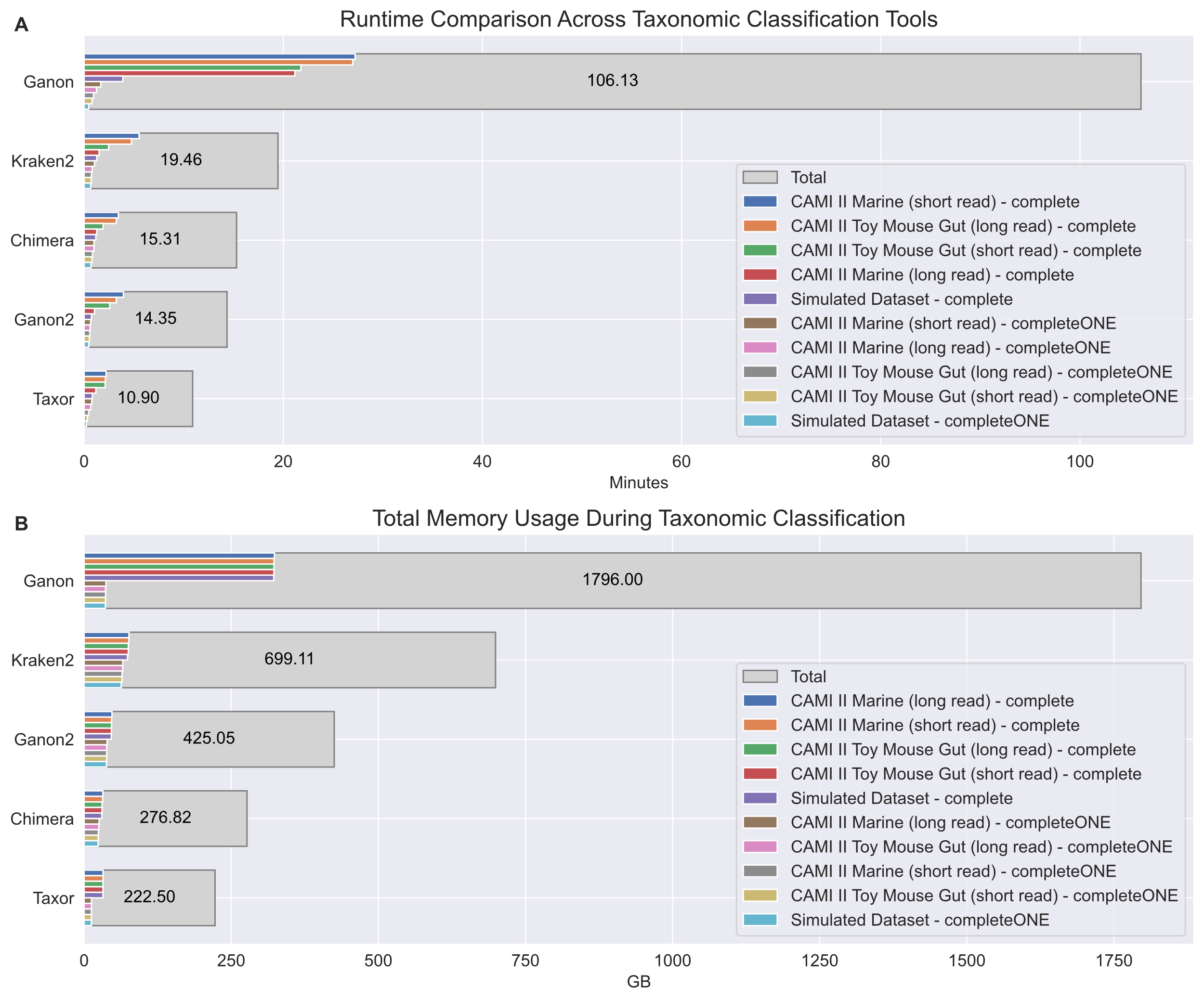

Figure S2 Runtime and Memory Usage of Taxonomic Classification Tools. (A) shows the runtime (in minutes), and (B) illustrates peak memory usage (in GB), the gray bars represent the total values across all datasets.

Table S1 Overview of Datasets Used for Database Construction

| **Dataset Name** | **Size (GB)** | **Total Sequences** | **Base Pairs (BP)** | **Assemblies** | **Species Count** |
| --- | --- | --- | --- | --- | --- |
| **Archaea**  **(2024.10.10)** | 1.6 | 1,138 | 1,725,129,521 | 601 | 468 |
| **CompleteONE**  **(2024.9.26)** | 50.6 | 36,811 | 52,322,440,925 | 25,091 | 25,129 |
| **Complete**  **(2024.10.7)** | 179.4 | 119,259 | 189,361,721,135 | 58,672 | 25,129 |
| **Refseq**  **(2024.10.16)** | 3,248.30 | 51,704,208 | 3,430,607,682,023 | 403,253 | 73,376 |

Table S2 Performance Comparison of Database Construction Tools

| **Software** | **Dataset Name** | **Build Time** | **Memory (GB)** | **Database Size (GB)** |
| --- | --- | --- | --- | --- |
| **Chimera** | Archaea (2024.10.10) | 7s | 0.9 | 0.4 |
| **Kraken2** |  | 6m30s | 26.7 | 2.1 |
| **Bracken** |  | 5m12s | 3.0 | 2.1+0.4 |
| **Ganon** |  | 23s | 2.9 | 0.6 |
| **Ganon2** |  | 53s | 3.4 | 1.4 |
| **Taxor** |  | 1m19s | 1.8 | 0.4 |
| **Chimera** | CompleteONE (2024.9.26) | 3m13s | 25.5 | 23.4 |
| **Kraken2** |  | 4h11m46s | 66.6 | 62.7 |
| **Bracken** |  | 6h41m55s | 66.2 | 62.7+36.7 |
| **Ganon** |  | 18m55s | 40.4 | 36.1 |
| **Ganon2** |  | 38m7s | 56.2 | 37.8 |
| **Taxor** |  | 3h56m22s | 18.6 | 12.2 |
| **Chimera** | Complete (2024.10.7) | 5m23s | 31.8 | 29.7 |
| **Kraken2** |  | 14h31m30s | 74.2 | 73.4 |
| **Bracken** |  | 9h15m7s | 76.6 | 73.4+60.7 |
| **Ganon** |  | 2h50m46s | 991.1 | 322.2 |
| **Ganon2** |  | 2h5m24s | 65.4 | 46.1 |
| **Taxor** |  | 6h30m18s | 38.5 | 32.1 |
| **Chimera** | Refseq (2024.10.16) | 2h1m53s | 170.9 | 168.6 |
| **Kraken2** |  | Failed due to memory overflow | | |
| **Bracken** |  |  |  |  |
| **Ganon** |  |  |  |  |
| **Ganon2** |  |  |  |  |
| **Taxor** |  |  |  |  |

Table S3 Overview of Sequencing Datasets for Taxonomic Classification Benchmarking

| **Dataset Name** | **Total Size (GB)** | **Samples** | **Read Length (bp)** | **Read Length S.D. (bp)** | **Insert Size Mean (bp)** | **Insert Size S.D. (bp)** |
| --- | --- | --- | --- | --- | --- | --- |
| **CAMI II Marine (long read)** | 10.1 | 10 | Average 3,000 | 1,000 | — | — |
| **CAMI II Marine (short read)** | 11.5 | 10 | 2 × 150 | — | 270 | 20 |
| **CAMI II Toy Mouse Gut (long read)** | 19.2 | 54 | Average 3,000 | 1,000 | — | — |
| **CAMI II Toy Mouse Gut (short read)** | 26.7 | 64 | 2 × 150 | — | 270 | 20 |
| **Simulated Dataset** | 18.4 | 480 | 2 × 100 or 2 × 150 | — | 200, 300, 400 | 10, 25, 50 |

### Supplemental Information

**Datasets for Database Construction**

All databases used in this study were constructed using NCBI RefSeq data, downloaded via Genome Updater (<https://github.com/pirovc/genome_updater>) to ensure consistency and up-to-date genomic information. To streamline the selection and processing of reference genomes, Chimera integrates an interactive Genome Updater module, allowing users to customize dataset downloads and automatically generate the Chimera-compatible target.tsv file, which is essential for database construction. This automation simplifies the process and ensures reproducibility across different database configurations.

To evaluate the efficiency and scalability of various database construction tools, we utilized multiple subsets of RefSeq data, each differing in size and complexity (see Table S1). The Archaea dataset includes all complete archaeal genomes from RefSeq and was selected to assess database construction performance on a relatively small dataset. The CompleteONE dataset consists of one complete genome per species, offering a representative yet compact reference that facilitates classification tool evaluations under reduced data volume constraints. Expanding upon this, the Complete dataset comprises all complete genomes available in RefSeq, enabling large-scale database construction scenarios to be tested, particularly regarding computational efficiency and storage demands. Finally, the RefSeq dataset includes all genomic data available in RefSeq, encompassing complete and draft genomes, viral sequences, plasmids, and other genomic elements. As the largest dataset used in this study, RefSeq serves as the ultimate benchmark for evaluating classification performance under extreme data scale conditions.

All datasets were retrieved using Genome Updater, and genomic sequences were stored in compressed fna.gz format to optimize storage efficiency. Since different classification tools require distinct input formats, a standardized preprocessing pipeline was applied to all datasets. Chimera employs an automated workflow that directly processes fna.gz files to generate the required target.tsv file, which is subsequently used for database construction.

**Chimera Database Construction**

Chimera (1.6.0) constructs databases using the chimera build command, with default parameters including a load factor of 0.58 and a maximum hash count of 2,000,000 per species, which are suitable for most datasets. The construction process relies on the target.tsv file generated by Chimera, which is compatible with Ganon’s target_info.tsv format, ensuring interoperability between tools.

For standard database construction, Chimera is executed with the following commands using its default settings:

```
# Construct the CompleteONE database
/usr/bin/time -v chimera build -i completeONE/target.tsv -o completeONEDB -t 32
# Construct the Complete database
/usr/bin/time -v chimera build -i complete/target.tsv -o completeDB -t 32
# Construct the RefSeq database
/usr/bin/time -v chimera build -i NCBIRefseq/target.tsv -o refseqDB -t 32
```

For smaller datasets, such as Archaea, increasing the load factor can optimize storage efficiency and reduce memory overhead. The following command constructs the Archaea database with a customized load factor of 0.95:

/usr/bin/time -v chimera build -i Archaea/target.tsv -t 32 -o ArchaeaDB --load-factor 0.95

**Kraken2 Database Construction**

Kraken2 v2.1.3 was used for database construction in this study. The construction process involved downloading the NCBI taxonomy database, decompressing genome sequence files, adding sequences to the Kraken2 library, and finally building the index. Since Kraken2 requires each genome sequence to be individually processed and added to the database, the time spent on adding sequences to the Kraken2 database is often comparable to or even longer than the index construction time. Notably, in the main text, the reported database construction time includes only the index-building step and does not account for the time required to add sequences to the database.

The following commands were used to construct the Archaea database:

```
# Download the NCBI taxonomy database
kraken2-build --download-taxonomy --db Archaea
# Copy genome sequence files to the Kraken2 database directory
find ~/tianqinzhong/project/chimera/archaea_files -type f -name '*.fna.gz' -exec cp {} ~/tianqinzhong/project/kraken2/Archaea/files/ \;
# Decompress all sequence files
find ~/tianqinzhong/project/kraken2/Archaea/files -name "*.gz" -print0 | xargs -0 -P32 -n1 gunzip
# Add sequence files to the Kraken2 database
find ~/tianqinzhong/project/kraken2/Archaea/files -name "*.fna" -print0 | xargs -0 -n1 -I {} kraken2-build --add-to-library {} --db Archaea
# Build the Kraken2 database
/usr/bin/time -v kraken2-build --build --db Archaea --threads 32
```

The following commands were used to construct the CompleteONE database:

kraken2-build --download-taxonomy --db CompleteONE

find ~/tianqinzhong/project/chimera/completeONE_files -type f -name '*.fna.gz' -exec cp {} ~/tianqinzhong/project/kraken2/CompleteONE/files/ \;

find ~/tianqinzhong/project/kraken2/CompleteONE/files -name "*.gz" -print0 | xargs -0 -P32 -n1 gunzip

find ~/tianqinzhong/project/kraken2/CompleteONE/files -name "*.fna" -print0 | xargs -0 -n1 -I {} kraken2-build --add-to-library {} --db CompleteONE

/usr/bin/time -v kraken2-build --build --db CompleteONE --threads 32

The following commands were used to construct the Complete database:

kraken2-build --download-taxonomy --db Complete

find ~/tianqinzhong/project/chimera/complete_files -type f -name '*.fna.gz' -exec cp {} ~/tianqinzhong/project/kraken2/Complete/files/ \;

find ~/tianqinzhong/project/kraken2/Complete/files -name "*.gz" -print0 | xargs -0 -P32 -n1 gunzip

find ~/tianqinzhong/project/kraken2/Complete/files -name "*.fna" -print0 | xargs -0 -n1 -I {} kraken2-build --add-to-library {} --db Complete

/usr/bin/time -v kraken2-build --build --db Complete --threads 32

The following commands were used to construct the RefSeq database:

kraken2-build --download-taxonomy --db RefSeq

find ~/tianqinzhong/project/chimera/refseq_files -type f -name '*.fna.gz' -exec cp {} ~/tianqinzhong/project/kraken2/RefSeq/files/ \;

find ~/tianqinzhong/project/kraken2/RefSeq/files -name "*.gz" -print0 | xargs -0 -P32 -n1 gunzip

find ~/tianqinzhong/project/kraken2/RefSeq/files -name "*.fna" -print0 | xargs -0 -n1 -I {} kraken2-build --add-to-library {} --db RefSeq

/usr/bin/time -v kraken2-build --build --db RefSeq --threads 32

**Bracken Database Construction**

In this study, Bracken v2.9 was used for taxonomic abundance estimation, leveraging pre-built Kraken2 databases to generate Bracken-compatible k-mer statistical models. Bracken improves species abundance estimation by reassigning Kraken2 classification results. Due to the large number of files being processed, it is necessary to first set the shell stack size to avoid the "parameter list too long" error. All parameters follow the default settings recommended by Bracken.

The following command constructs the Archaea database:

```
# Set shell stack size
ulimit -s 1024000
# Build the Bracken database
/usr/bin/time -v bracken-build -d Archaea -t 32 -k 35 -l 100
```

The commands for constructing other databases (CompleteONE, Complete, and RefSeq) are as follows:

```
# Build the CompleteONE database
/usr/bin/time -v bracken-build -d CompleteONE -t 32 -k 35 -l 100
# Build the Complete database
/usr/bin/time -v bracken-build -d Complete -t 32 -k 35 -l 100
# Build the RefSeq database
/usr/bin/time -v bracken-build -d RefSeq -t 32 -k 35 -l 100
```

**Ganon and Ganon2 Database Construction**

The database construction process for Ganon and Ganon2 (2.1.0) is identical, except that Ganon enables Interleaved Bloom Filters (IBF) using the -v ibf option, while Ganon2 does not use this parameter.

Below are the commands for constructing the Archaea, CompleteONE, Complete, and RefSeq databases:

\# Construct Archaea database (Ganon)

/usr/bin/time -v ganon build --source refseq --organism-group archaea --threads 32 --complete-genomes --db-prefix Archaea -v ibf

\# Construct Archaea database (Ganon2)

/usr/bin/time -v ganon build --source refseq --organism-group archaea --threads 32 --complete-genomes --db-prefix Archaea

\# Construct CompleteONE database (Ganon)

/usr/bin/time -v ganon build --source refseq --organism-group archaea bacteria fungi viral --threads 32 --complete-genomes --genome-updater "-A 'species:1'" --db-prefix CompleteONE -v ibf

\# Construct CompleteONE database (Ganon2)

/usr/bin/time -v ganon build --source refseq --organism-group archaea bacteria fungi viral --threads 32 --complete-genomes --genome-updater "-A 'species:1'" --db-prefix CompleteONE

\# Construct Complete database (Ganon)

/usr/bin/time -v ganon build --source refseq --organism-group archaea bacteria fungi viral --threads 32 --complete-genomes --db-prefix Complete -v ibf

\# Construct Complete database (Ganon2)

/usr/bin/time -v ganon build --source refseq --organism-group archaea bacteria fungi viral --threads 32 --complete-genomes --db-prefix Complete

\# Construct RefSeq database (Ganon)

/usr/bin/time -v ganon build --source refseq --threads 32 --db-prefix RefSeq --organism-group archaea bacteria fungi human invertebrate metagenomes other plant protozoa vertebrate_mammalian vertebrate_other viral -v ibf

\# Construct RefSeq database (Ganon2)

/usr/bin/time -v ganon build --source refseq --threads 32 --db-prefix RefSeq --organism-group archaea bacteria fungi human invertebrate metagenomes other plant protozoa vertebrate_mammalian vertebrate_other viral

**Taxor Database Construction**

This study utilizes Taxor v0.1.3 for database construction. The process involves taxonomy database preparation, taxonomic information extraction, and index construction. All sequence files must be stored in a single directory without subfolders; otherwise, taxor build will not process them correctly. Additionally, when using Genome Updater to download data, the -a option can be added to ensure the taxonomy database remains up to date.

Taxonomy Database Preparation

mkdir -p taxdump

tar -zxvf taxdump.tar.gz -C taxdump

Generating Taxonomic Information File

cut -f 1,7,20 assembly_summary.txt \

| taxonkit lineage -i 2 -r -n -L --data-dir taxdump \

| taxonkit reformat -I 2 -P -t --data-dir taxdump \

| cut -f 1,2,3,4,6,7 > refseq_accessions_taxonomy.csv

Database Index Construction

```
# Construct Archaea Database
/usr/bin/time -v taxor build --input-file archaea.csv \
--input-sequence-dir archaea/files \
--output-filename archaea.hixf \
--threads 32 --kmer-size 22 --syncmer-size 12 --use-syncmer
# Construct CompleteONE Database
/usr/bin/time -v taxor build --input-file completeONE.csv \
--input-sequence-dir completeONE/2024-09-26_11-57-14/files \
--output-filename completeONE.hixf \
--threads 32 --kmer-size 22 --syncmer-size 12 --use-syncmer
# Construct Complete Database
/usr/bin/time -v taxor build --input-file complete.csv \
--input-sequence-dir complete/files \
--output-filename complete.hixf \
--threads 32 --kmer-size 22 --syncmer-size 12 --use-syncmer
# Construct RefSeq Database
/usr/bin/time -v taxor build --input-file refseq.csv \
--input-sequence-dir refseq/files \
--output-filename refseq.hixf \
--threads 32 --kmer-size 22 --syncmer-size 12 --use-syncmer
```

**Taxonomic Classification Tools and Execution**

This study employs Chimera, Kraken2, Bracken, Ganon, Ganon2, and Taxor for taxonomic classification experiments to evaluate the performance of different classification tools on metagenomic datasets. All tools are executed with their officially recommended parameters and evaluated on multiple datasets.

All tools run in the same computational environment, using the same number of threads ({threads}) and classification thresholds ({threshold}) to ensure fairness in benchmarking.

Classification Execution

```
# Chimera
conda activate chimera
/usr/bin/time -v chimera classify -i {seq_path} -d {db_path}DB.imcf -t {threads} -s {threshold} -o {output}
# Ganon
conda activate ganon
/usr/bin/time -v ganon classify -d {db_path} -s {seq_path} -t {threads} -c {threshold} -o {output} --verbose --output-all --output-one
# Ganon2
conda activate ganon
/usr/bin/time -v ganon classify -d {db_path} -s {seq_path} -t {threads} -c {threshold} -o {output} --verbose --output-all --output-one
# Kraken2
conda activate kraken2
/usr/bin/time -v kraken2 --db {db_path} --threads {threads} --confidence {threshold} --output {output} {extra} {seq_path} --report {report}  # Report file required for Bracken
# Taxor
conda activate taxor
/usr/bin/time -v taxor search --index-file {db_path}.hixf --query-file {seq_path} --output-file {output} --percentage {threshold} --threads {threads}
# Bracken
conda activate kraken2
/usr/bin/time -v bracken -d {db_path} -i {report} -o {bracken_output}
```

**Performance Evaluation Metrics**

This study employs multiple performance evaluation metrics to comprehensively assess the effectiveness of different taxonomic classification tools on metagenomic data. All evaluations are based on the comparison between classification results and reference taxonomic annotation files.

Evaluation Metrics

Accuracy: Defined as the proportion of correctly classified instances, calculated as:

$$Accuracy=\frac{TP+TN}{TP+TN+FP+FN}$$

Precision: Measures the accuracy of the classifier when predicting positive categories, calculated as:

$$Precision=\frac{TP}{TP+FP}$$

Recall (Sensitivity): Represents the proportion of actual positive instances correctly identified by the classifier, calculated as:

$$Recall=\frac{TP}{TP+FN}$$

F1 Score: The harmonic mean of precision and recall, balancing their effects, calculated as:

$$F1=2\times\frac{Precision\times Recall}{Precision+Recall}$$

L1 Distance: Measures the deviation between classification results and the true taxonomic distribution, calculated as:

$$L1=\sum_{i} |P_{i}-T_{i}|$$

where $P_{i}$ represents the normalized abundance of the predicted category, and $T_{i}$ represents the normalized abundance of the true category.

Classification Result Definitions

TP (True Positive): DNA sequences correctly classified into their species.

FN (False Negative): DNA sequences that failed to be classified.

FP (False Positive): DNA sequences misclassified as another species.

TN (True Negative): DNA sequences correctly classified as non-target species.

**Automated Classification and Result Compilation Pipeline**

This study developed an automated classification and result compilation pipeline to ensure that different classification tools run under identical experimental conditions while standardizing classification results and performance evaluation. This pipeline is implemented using a Python automation script (<https://github.com/LoadStar822/ChimeraBenchmark/blob/master/runClassifier.py>), enabling batch execution of Chimera, Kraken2, Bracken, Ganon, Ganon2, and Taxor, along with automatic result processing.

Due to the large number of files requiring classification, we further developed a command generation script (<https://github.com/LoadStar822/ChimeraBenchmark/blob/master/generate_and_run.py>), which automatically generates execution commands for each tool and batch-submits them to runClassifier.py, thereby improving task scheduling efficiency.

The classification pipeline consists of automated classification execution, result formatting, and performance evaluation based on reference taxonomic annotation files. First, execution parameters for all classification tools, including database paths, thread counts, and classification thresholds, are preconfigured to ensure all tools operate under the same computational environment for fair comparisons. Then, the command generation script generate_and_run.py creates classification commands for all tools and submits them to runClassifier.py for automated execution.

Subsequently, the script compares classification results against reference taxonomic annotation files (ground truth) and calculates classification performance metrics, including Accuracy, Precision, Recall, F1 Score, and L1 Distance. Finally, a CSV file containing classification performance results for all tools is generated.

**Database Construction with FMC**

Chimera provides the FMC strategy, which controls the maximum number of minimizer hashes per species using the --max-hashes parameter. This reduces taxonomic overrepresentation and optimizes database storage. Setting --max-hashes to 0 disables FMC.

**FMC Integration into Ganon**

To evaluate the generalizability of the FMC strategy, we integrated it into the Ganon classification tool and conducted experiments on the same datasets to assess its impact on classification performance and resource utilization.

We modified Ganon's source code by adjusting the count_hashes function in the GanonBuild.cpp file, enforcing a maximum minimizer hash limit of 2000000 during database construction. The modified version was then recompiled and executed.

Experimental results demonstrate that FMC is equally effective in Ganon, optimizing database structure and enhancing classification performance, proving that this strategy can be extended to various metagenomic classification tools.
